# Topologically Associating Domain Boundaries are Commonly Required for Normal Genome Function

**DOI:** 10.1101/2021.05.06.443037

**Authors:** Sudha Rajderkar, Iros Barozzi, Yiwen Zhu, Rong Hu, Yanxiao Zhang, Bin Li, Yoko Fukuda-Yuzawa, Guy Kelman, Adyam Akeza, Matthew J. Blow, Quan Pham, Anne N. Harrington, Janeth Godoy, Eman M. Meky, Kianna von Maydell, Catherine S. Novak, Ingrid Plajzer-Frick, Veena Afzal, Stella Tran, Michael E. Talkowski, K. C. Kent Lloyd, Bing Ren, Diane E. Dickel, Axel Visel, Len A. Pennacchio

**Author notes:** To whom correspondence should be addressed: D.E.D.,; A.V.,; L.A.P.

## Abstract

Topologically associating domain (TAD) boundaries are thought to partition the genome into distinct regulatory territories. Anecdotal evidence suggests that their disruption may interfere with normal gene expression and cause disease phenotype^1–3^, but the overall extent to which this occurs remains unknown. Here we show that TAD boundary deletions commonly disrupt normal genome function *in vivo*. We used CRISPR genome editing in mice to individually delete eight TAD boundaries (11-80kb in size) from the genome in mice. All deletions examined resulted in at least one detectable molecular or organismal phenotype, which included altered chromatin interactions or gene expression, reduced viability, and anatomical phenotypes. For 5 of 8 (62%) loci examined, boundary deletions were associated with increased embryonic lethality or other developmental phenotypes. For example, a TAD boundary deletion near *Smad3/Smad6* caused complete embryonic lethality, while a deletion near *Tbx5/Lhx5* resulted in a severe lung malformation. Our findings demonstrate the importance of TAD boundary sequences for *in vivo* genome function and suggest that noncoding deletions affecting TAD boundaries should be carefully considered for potential pathogenicity in clinical genetics screening.

## Introduction

Eukaryotic genomes fold into topologically associating domains (TADs), sub-megabase-scale chromatin segments characterized by high intra-domain chromatin contact frequency^4–6^. TADs represent a key feature of hierarchical genome organization by defining chromatin neighborhoods within which regulatory sequences can interact, while simultaneously insulating regulatory interactions across boundaries^5,7–10^. TAD boundaries are primarily defined and measured through chromatin conformation assays, and they are typically associated with a signature set of proteins, including CCCTC-binding factor (CTCF), proteins in the structural maintenance of chromosomes (SMC) complex such as cohesin and condensin, and RNA polymerase II^5,11–14^. TADs form as a result of loop extrusion, wherein DNA strands slide from within the cohesin or SMC complex until bound CTCF molecules in a convergent orientation are met^13,15–19^. Loss of CTCF, cohesin, or the cohesin loading factor Nipbl results in TAD disruption, while loss of cohesin release factor, Wapl, results in reinforcement of TAD boundaries^20,21^. Intriguingly, approximately 20% of TAD boundaries remain stable upon loss of CTCF^22^. Both CTCF-mediated mechanisms and transcription can affect the formation and function of TADs, but neither seems to be individually sufficient nor universally required^7,11,23,24^. Thus, chromatin state, transcriptional activity, and TAD organization may influence each other, and the observed nuclear structure of mammalian genomes likely results from their complex interplay^7,11,23–25^.

The genomic locations of TAD boundaries are well conserved across mammalian species, suggesting that their function and positions within the genome are subject to evolutionary constraint^5,26–28^. This notion is further supported by the overall depletion of structural variants at TAD boundaries observed in the general human population^26^, while disruptions and rearrangements of TAD structure have been implicated in the mis-expression of genes and are associated with developmental and cancer phenotypes^1–3, 29, 30^,. However, most of these disruptions were spontaneously occurring structural mutations that also included neighboring genomic features, such as regulatory elements and/or protein-coding genes. Therefore, the specific role of TAD boundaries in these phenotypes is not well understood. In the present study, we examine the generalized functional necessity of TAD boundary sequences *in vivo*. We selected eight independent TAD boundaries in the vicinity of genes active during embryonic development, individually deleted these boundaries from the mouse genome, and systematically examined the consequences on survival, genome organization, gene expression, and development. All eight TAD boundary deletions caused alterations of at least one of these properties. We also observed that loss of boundaries with more CTCF sites generally resulted in more severe phenotypes and that both organismal and regulatory phenotypes coincided with marked changes in chromatin conformation in at least one TAD boundary deletion. In combination, our results indicate that TAD boundary sequences are commonly required for normal genome function and development.

## Results

### Strategy for selecting TAD boundaries for *in vivo* deletion

Presence of CTCFs, along with the cohesin complex proteins, is the hallmark of TAD boundaries that are conserved across cell types in closely associated species^13,15–19^. In order to assess the *in vivo* functions of TAD boundary sequences, we prioritized boundaries with these prototypical features and, in addition, we focused on boundaries of TADs that contain genes with known expression and function in embryonic development. We hypothesized that selecting TAD boundaries with flanking TADs that harbored genes with known developmental phenotypes would increase the likelihood of detecting organismal phenotypes and guide the phenotyping of boundary deletion mice. From a genome-wide set of >3,300 annotated TAD boundaries^5,10^, we scored and prioritized each boundary based on the following critical criteria: 1) CTCF occupancy aggregated from 62 published CTCF ChIP-seq datasets, which served as a proxy for the expected overall strength of insulation (datasets listed in **Supplementary Table 1**); 2) co-occupancy of subunits of the cohesin complex and the transcription factor Znf143^31^ from 38 published ChIP-seq datasets (**Supplementary Table 1**); 3) CTCF-binding conservation at orthologous regions in four different mammalian species^32^ (**Supplementary Table 2**); and 4) whether both flanking TADs contain genes with known roles in embryonic development, preferentially showing divergent patterns of tissue-specific expression (**Figure 1, Extended Data Figure 1, Supplementary Tables 1-3, Methods**). TAD boundaries that encompassed protein-coding genes were excluded. Following genome-wide prioritization, we selected and deleted 8 individual TAD boundaries from the mouse genome through pronuclear injection of fertilized eggs using CRISPR/Cas9. These deletions ranged in size from 11 to 80 kb and removed all known CTCF and cohesin binding sites in each TAD boundary region, while leaving any nearby protein-coding genes intact (**Figure 1, Extended Data Figures 1-2, Supplementary Table 3, Methods**). Live founder mice heterozygous for the targeted deletion were successfully obtained for all eight boundaries, and these were bred into stable lines to assess molecular and organismal phenotypes.

**Figure 1.**
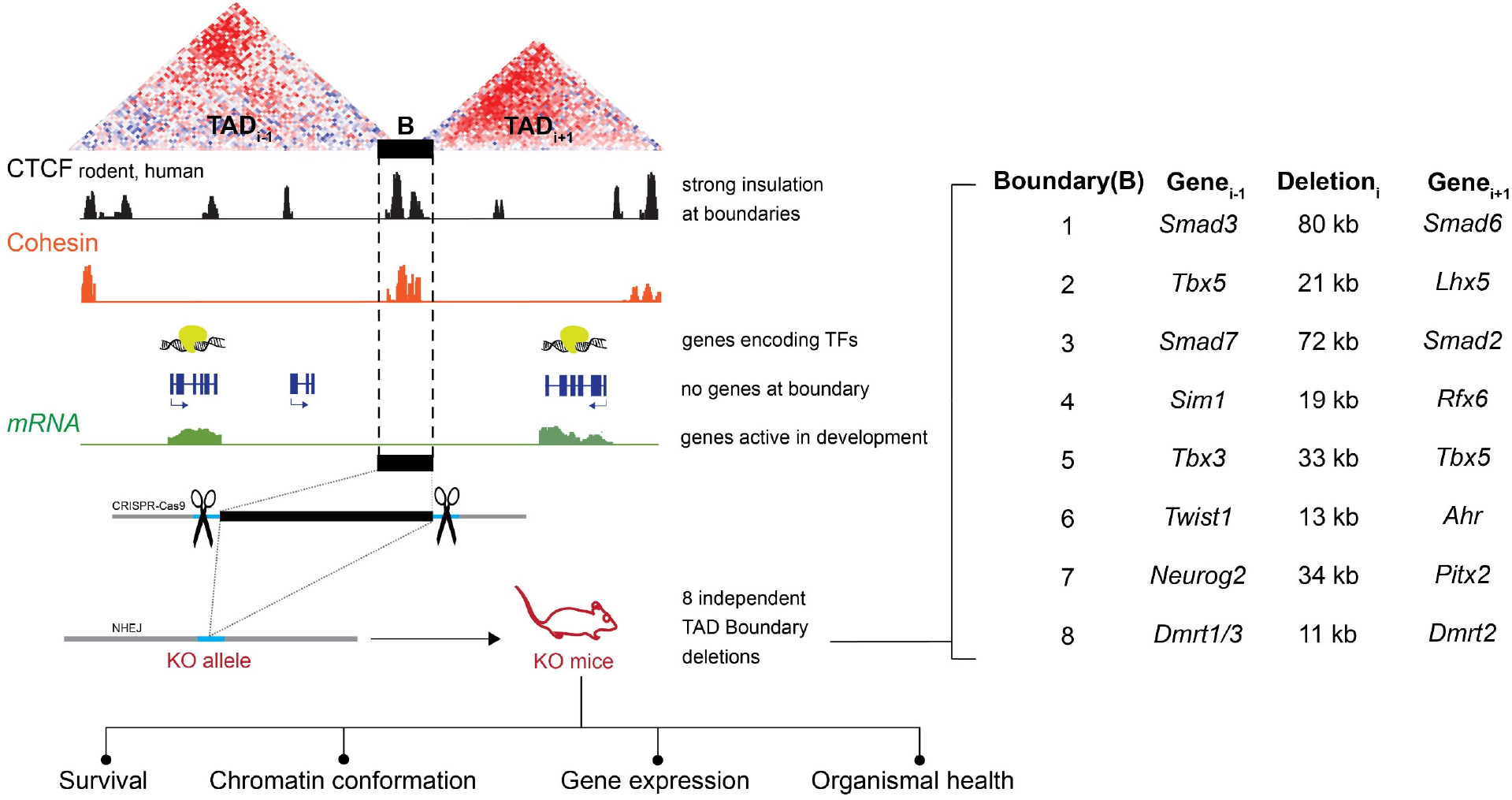
Study overview. **Left:** Schematic showing the selection and CRISPR/Cas9-based deletion strategy for removal of TAD boundaries *in vivo*, along with types of phenotyping performed on the resulting knock-out (KO) mice. **Right:** the specific boundaries individually deleted, along with selected developmentally expressed genes flanking each deleted boundary. B, boundary element; TFs, transcription factors.

### TAD boundary deletions disrupt prenatal development

To investigate the *in vivo* consequences of TAD boundary deletions, we assessed all lines for viability in homozygous offspring (**Figure 2a**). First, we cross-bred heterozygous deletion mice to determine if homozygous offspring were viable and present at the Mendelian-expected rate (25%). For line B1, in which a boundary between *Smad3* and *Smad6* had been deleted, no mice homozygous for the deletion were obtained out of 329 live births. Timed breeding revealed that homozygous embryos are present at embryonic day 8.5 (E8.5) at the expected Mendelian ratio but not at later stages of development (p<0.05, Chi-squared test, for all examined stages E10.5 and later; **Figure 2b**). While no viable homozygous-null embryos were observed at E10.5, we observed partially resorbed homozygous deletion embryos at this stage, further corroborating that homozygous deletion of boundary B1 cause fully penetrant loss of viability between E8.5 and E10.5 (**Figure 2c**). Homozygous deletion of four additional boundary loci were associated with partially penetrant embryonic or perinatal lethality (**Figure 2a**). For the most extreme (B2), we observed a loss of approximately 65% of expected homozygous offspring at weaning (p=3.90E-10, Chi-squared test; **Figure 2a, Extended Data Figure 2**). For three additional boundary deletions (B3, B4, B5), we observed a 20-37% depletion of homozygotes at weaning (p<0.05, Chi-squared test, in all cases, **Figure 2a**, **Extended Data Figure 2**). There were no significant sex biases among viable homozygous offspring in any of the lines (**Extended Data Figure 3**). Overall, these data identify several examples of individual boundary elements that are required for normal organismal viability.

**Figure 2.**
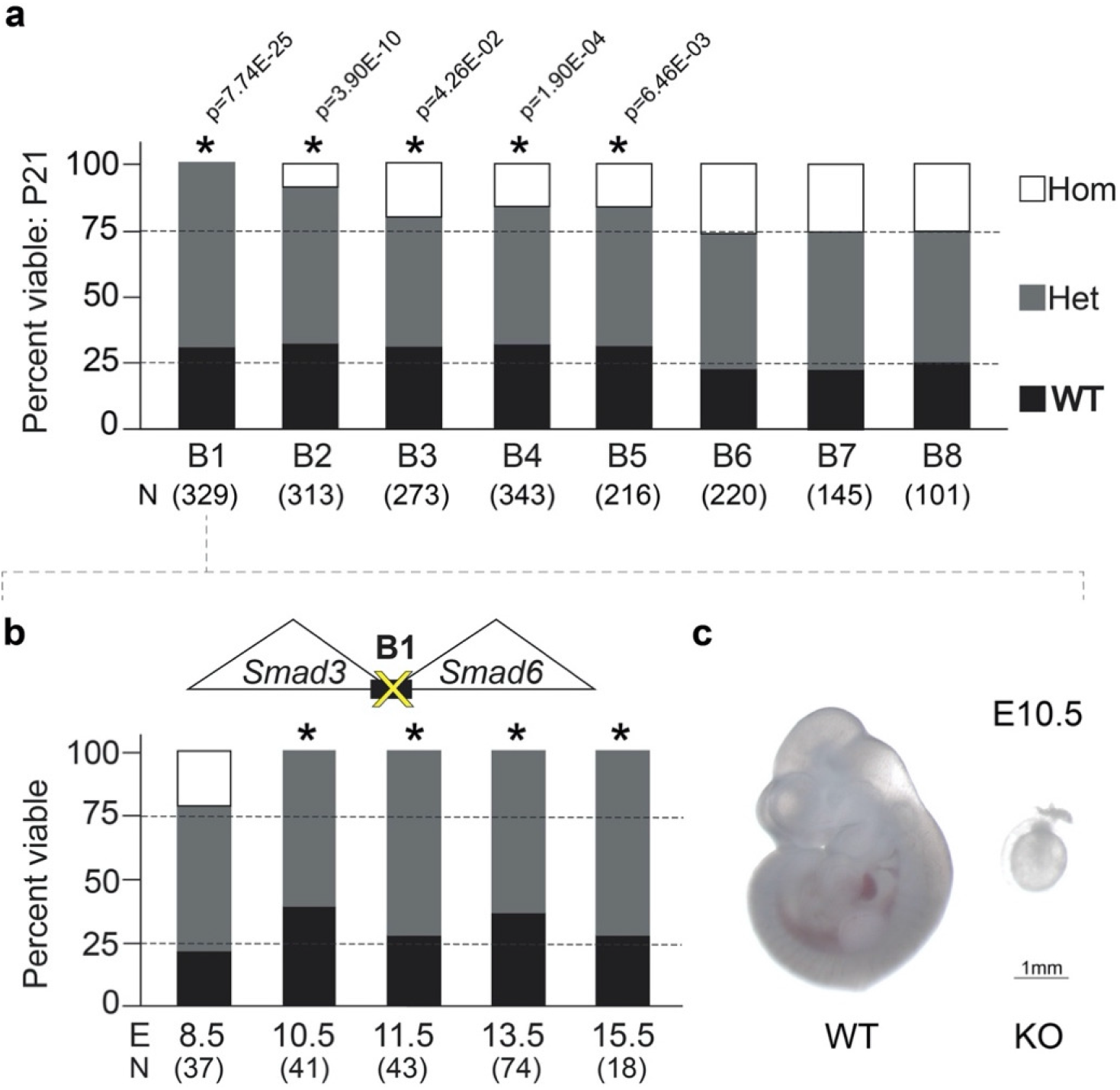
TAD boundary deletions result in reduced viability. **a.** Mendelian segregation of offspring from heterozygous crosses at weaning for all TAD boundary deletion lines. P21, postnatal day 21; N, number of pups analyzed; Hom, homozygous; Het, heterozygous; WT, wild-type. **b.** Mendelian segregation of offspring from heterozygous crosses at designated embryonic stages for boundary deletion locus B1 (* indicates p<0.05). E, embryonic day; N, number of embryos analyzed. **c.** Brightfield images of representative littermate wild-type (left) and homozygous mutant (right) embryos obtained at E10.5. Scale bar, 1mm.

### TAD boundary deletions result in abnormal TAD architecture

To assess the effects of TAD boundary deletions on the chromatin interaction landscape, we performed high-throughput chromosome conformation capture (i.e., Hi-C) in TAD boundary knockout mice and wild-type controls for the seven lines where viable homozygous deletion mice were obtained (**Figure 3, Extended Data Figure 5**). For three loci, we observed that homozygous-null mice displayed loss of insulation at the TAD boundaries and concurrent merging of neighboring TADs (loci B2, B3, and B6; **Figure 3**). Loss of insulation was also observed at a fourth TAD boundary deletion (B8), but this was not associated with major changes to the overall TAD configuration at this locus (**Extended Data Figure 4**). As a second measure of disrupted chromatin structure, we compared the directionality index (DI) between knockout mice and wildtype controls. DI assesses the trend of upstream (leftward or negative) or downstream (rightward or positive) contacts along a region of the chromosome^5^ and corner regions peripheral to TADs where abrupt shifts in upstream and downstream contacts are observed (i.e., sites with statistically significant contact biases are computationally called as boundaries between flanking TADs). Changes in DI were observed in homozygous-null mutants in 5 of the 7 lines assessed by Hi-C (p≤0.01, Wilcoxon rank-sum test, for loci B2, B3, B4, B6, and B8; **Figure 3**, **Extended Data Figure 4**). Moreover, for three TAD boundary deletions (loci B4, B7, and B8) we observed a reduction of long-range contacts in one of the TADs adjacent to the deleted boundary (**Extended Data Figure 4**). Taken together, we observed chromatin conformation changes in 86% (6/7) of lines examined. These data indicate that removal of individual TAD boundary sequences commonly affects insulation between neighboring TADs, resulting in altered interaction frequency between sequences in normally isolated domains.

**Figure 3.**
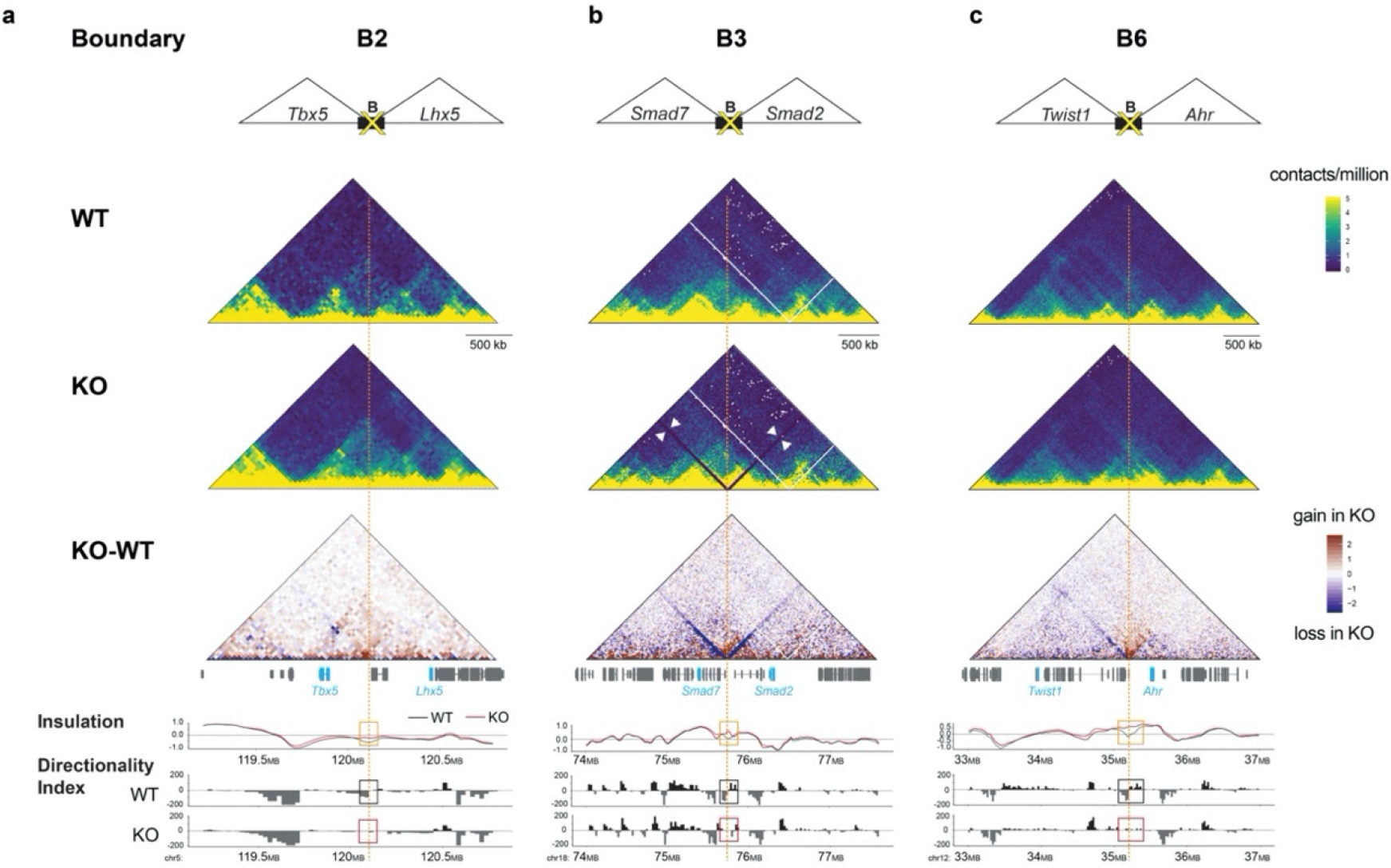
Boundary deletions result in abnormal TAD architecture. Hi-C derived interaction maps for TAD boundary loci B2 (**a**), B3 (**b**), and B6 (**c**). **Top**: cartoon at the top depicts the TAD boundary deleted, with select developmental genes within the TADs flanking the deleted boundary (B). **Middle:** Three heatmaps showing Hi-C contact data. The top two heat maps (yellow-blue color-code) show Hi-C contact matrices presented as observed/expected contacts at 25 kb resolution in representative wild-type (WT, top) and knockout (KO, middle) mouse liver tissue samples. The third heatmap (brown-blue color-code) shows net changes in Hi-C interaction frequencies in the KO relative to WT. Positions of genes within the corresponding locus are indicated below the heatmap. The dashed orange vertical line indicates the position of the deleted boundary. **Bottom:** The line graph below plots the insulation profiles for the WT and KO samples shown in the heat maps above. The insulation profile assigns an insulation score to each genomic interval^50^, with local minima representing the most insulated region. Note the deviation from the minima in the insulation profile for the KO compared to WT (orange box), indicating loss of insulation in the KO. The lower-most bar plots show the Directionality Index (DI) for samples shown in the plots above. DI measures the trend of upstream (leftward or negative) or downstream (rightward or positive) contacts at a given locus^5^. Boundaries are called at regions where abrupt and significant shifts in upstream and downstream contacts are observed, as depicted in the WT (black box). Note the loss of demarcation between upstream and downstream contacts at the deleted site in the KO (red box). White lines in the top two heat maps for boundary B3: artefact in mapping and/or alignment. Oxford blue lines (white arrowheads) in the heatmap for KO in B3: the 72 kb deletion of B3. More details on replicates and additional TAD boundary deletions are provided in **Extended Data Figure 5** and **Methods**.

### TAD boundary deletions cause molecular and developmental phenotypes

We next examined whether loss of TAD boundaries and resulting changes in chromatin architecture were associated with additional molecular or physiological phenotypes. To determine if the deletions altered the expression of genes in the vicinity of each TAD boundary, we measured gene expression in E11.5 embryos with homozygous boundary deletions and matched wild-type controls. For each line, RNA-seq was performed in two different tissues with known expression of the genes located in the adjacent TADs, and qPCR was performed to query select genes in a larger panel of tissues for a subset of the TAD boundary deletion lines (**Extended Data Figures 5-6**). Across all seven lines examined by this approach, we identified two cases in which expression of a gene(s) in a TADs flanking the boundary changed. Embryos homozygous-null for B2 displayed a 44% reduction in *Tbx5* in the lungs as compared to WT controls (p_adj_=0.04, **Extended Data Figure 4, Supplementary Table 4**). Embryos homozygous-null for B6 showed ~40% reduction of expression of three genes (*Meox2, Sostdc1*, and *Prkar2b* in the heart; *Meox2* in the developing face; p_adj_=0.04, **Extended Data Figure 5 and Supplementary Table 4**). These data show that deletion of TAD boundaries alone can result in pronounced changes in tissuespecific gene expression.

Since most TAD boundary deletions did not result in fully penetrant embryonic lethality, we comprehensively assessed postnatal phenotypes in all lines with viable homozygous offspring. We used a standardized panel of general anatomical, histological and necropsy examinations, including 230 sensory, neurological and behavioral tests, 20 tests measuring cardiac function, 35 metabolic function tests, 20 hematological/immunological parameters, and 36 musculo-skeletal tests, performed in 8-12 sex- and age-matched pairs of control and homozygous knockout mice from each line (>6000 data points across 350 total measurements; **Figure 4a, Extended Data Figure 7** and **Supplementary Table 5**). The most remarkable phenotype observed occurred in mice homozygous for deletions of boundary B2, located between *Tbx5* and *Lhx5*. As described above, mice homozygous-null for B2 were born at significantly reduced frequency. Surviving knockout mice showed severely underdeveloped lungs, with a vestigial left side (**Figure 4b**). This phenotype is partially penetrant (12 out of 20 mice, or 60%) with higher rates observed in male homozygous mutants (82%) than females (33%). This lung anomaly is consistent with the downregulation of the nearby *Tbx5* gene in the developing lungs, as the phenotype has been observed in lung-specific knockouts of *Tbx5* ^33^. This indicates that the deletion of boundary sequences can be sufficient to cause local gene dysregulation underlying severe phenotypes. Upon closer observation of the B2 boundary region, we identified a subregion showing lung-specific putative enhancer marks including open chromatin (ATAC-seq) and H3K27ac in a publicly available ENCODE atlas of regulatory regions^34,35^ (**Extended Data Figure 8a**). This subregion with enhancer-associated features is immediately adjacent to the primary CTCF site and was therefore included in the B2 boundary deletion interval (**Extended Data Figure 8a**). In a transgenic mouse reporter assay, this 852 bp subregion was sufficient to drive *lacZ* expression in lungs across multiple mouse embryonic stages. (**Extended Data Figure 8b and Methods**). These results suggest that molecular functions beyond CTCF-mediated insulation of chromatin domains may be part of TAD boundary regions. The enhancer activity observed in reporter assays is consistent with the observed gene expression changes in *Tbx5* and the accompanying lung defect. A genome-wide survey of the location of known embryonically active enhancers^36^ relative to predicted TAD boundaries^10^ suggests that other TAD boundaries across the genome contain enhancers that are part of or in the immediate vicinity (<5kb) of the boundary region (**Extended Data Figure 9**).

**Figure 4.**
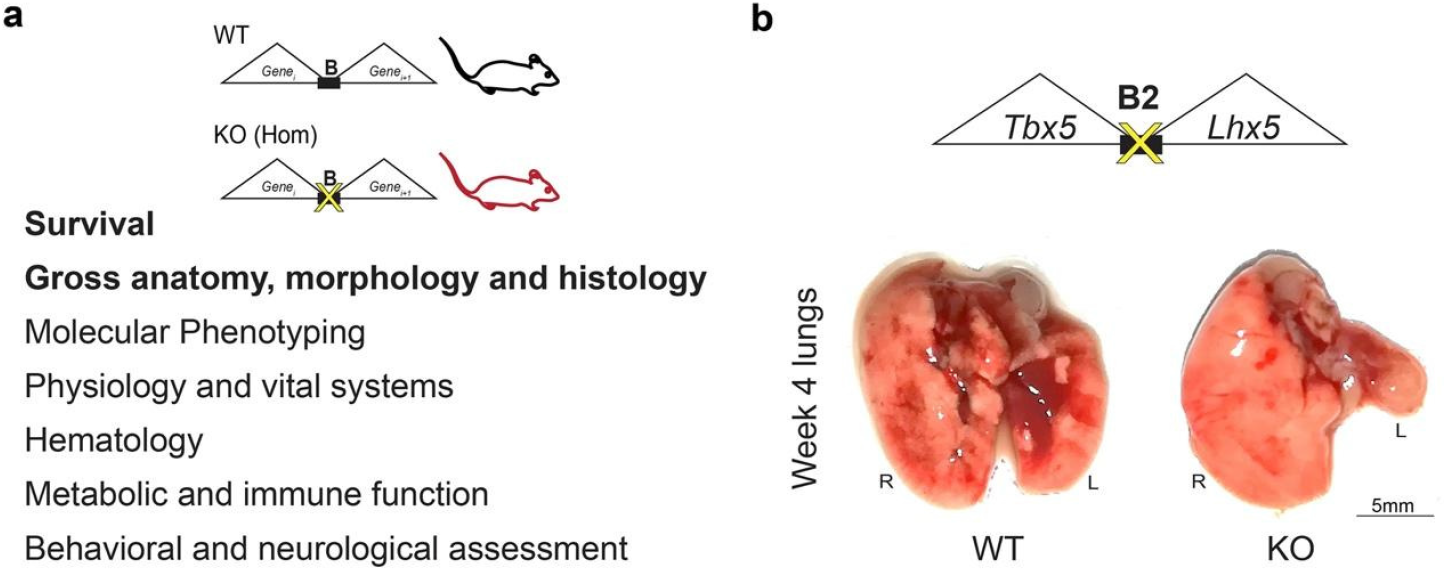
TAD boundary deletions result in developmental phenotypes. **a.** Overview of comprehensive phenotypic assessment performed to characterize effects of TAD boundary deletion *in vivo*. **b.** Homozygous mutants (KO) for TAD boundary deletion locus B2 show vestigial left lung. Scale bar, 5mm.

While our viability statistics suggest that other boundary deletions likely also impact *in vivo* biology, no overt phenotypes, or none with a clear link to the genes in the vicinity of the boundary, were observed for 6 out of 7 TAD boundary deletions despite extensive screening from birth through postnatal week 16 (**Extended Data Figure 7** and **Supplementary Table 5**). As we selected boundaries for those with flanking genes expressed during development, phenotypes may only be observable in embryonic or perinatal stages, and any surviving mice may not show major phenotypes after birth.

## Discussion

TAD boundaries have been hypothesized to be critical for normal genome function based on their known molecular roles in defining regulatory territories along chromosomes and in preventing enhancer-promoter interactions between adjacent chromatin domains. This notion was supported by observations of disease phenotypes associated with structural mutations that include TAD boundaries, although in most cases in combination with adjacent genomic features such as regulatory sequences or protein-coding genes^2,8,37,38^. To assess their general requirement for normal genome and organism function, we performed targeted genomic deletions of TAD boundaries in mice. Our studies focused on genome regions that included genes with known developmental roles in order to facilitate the detection of possible phenotypes. Remarkably, all eight TAD boundary deletions in engineered mice resulted in abnormal molecular or organismal phenotypes (summarized in **Figure 5**). This included six lines showing alterations to chromatin interactions within or across neighboring TADs, two lines with substantial alterations of expression level of neighboring gene(s), five lines displaying complete or partial embryonic lethality, and one line showing defects in lung development. Notably, mice with a deletion of boundary B2 display lung abnormalities that recapitulate a phenotype observed in mice lacking expression of the nearby *Tbx5* gene in the lung^33^, and we observed a sequence that is sufficient to drive gene expression in the lung directly adjacent to the primary CTCF site within the boundary sequence itself (**Extended Data Figure 8**). Taken together, our results support that TAD boundaries are commonly required for normal genome function and removal of the boundary region itself commonly causes alterations in chromatin structure, gene expression, or organism development. The boundary deletions performed in this study caused a spectrum of phenotypes, ranging from severe (complete embryonic lethality) to mild (molecular phenotypes only). Likewise, the chosen boundaries ranged widely in the number of CTCF clusters present, from two in boundaries that caused altered chromatin interactions or gene expression changes only, to three to five in the boundaries that cause reduced viability, to 11 CTCF clusters in the boundary that caused the most severe phenotype. The dosage of CTCF at TAD boundaries is known to affect the formation of TADs^22,39,40^, and while the number of boundaries studied here is too small to establish a statistically robust correlation, it is tempting to speculate that deletions of boundaries with more CTCF sites tend to cause more pronounced phenotypes.

**Figure 5.**
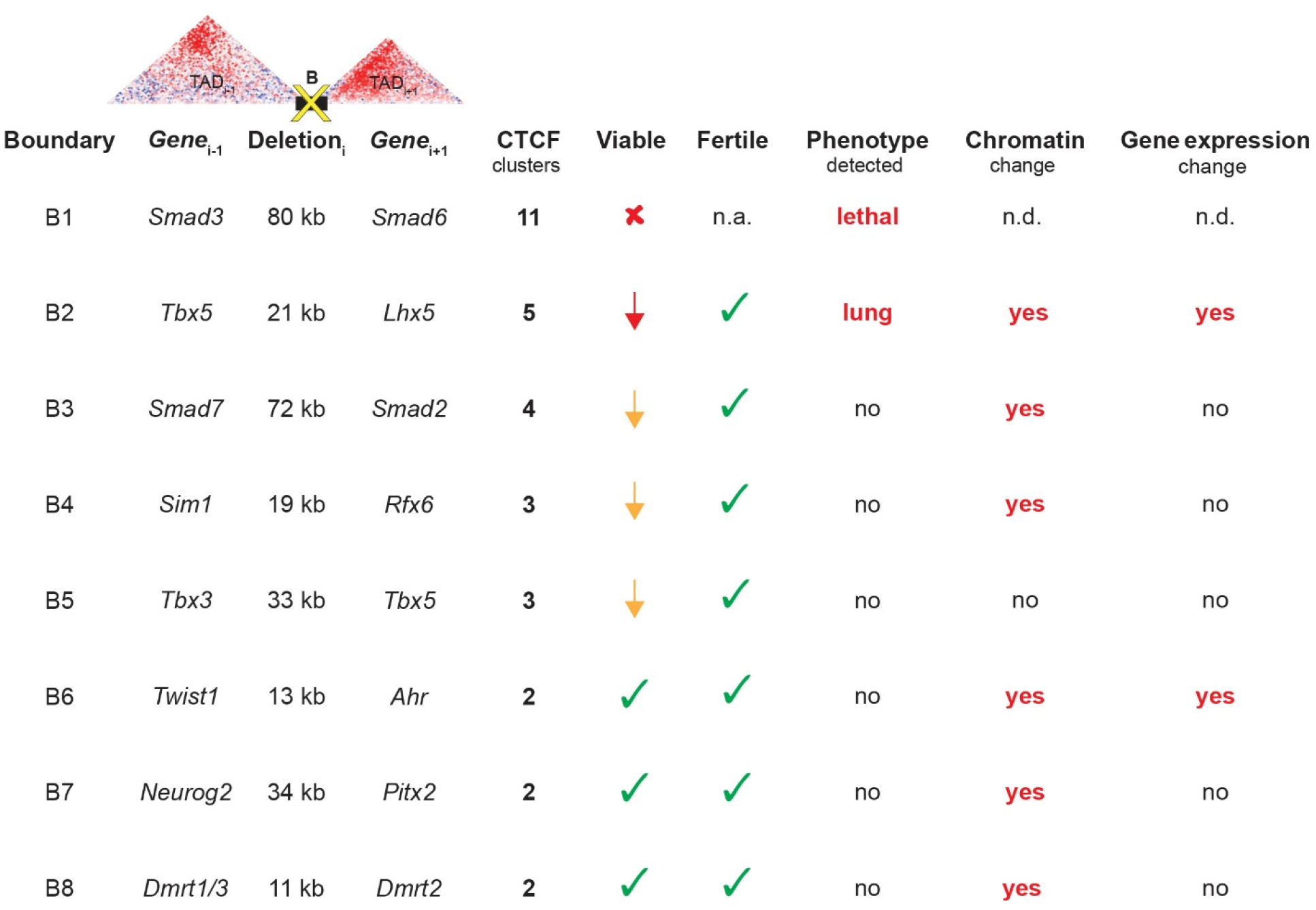
Summary of findings. Columns in the left show specific TAD boundaries (B1-8) deleted in this study, key developmental genes within the flanking TADs (Gene_i-1_ and Gene_i+1_), approximate boundary deletion sizes (Deletion_i_), and the number of putative CTCF clusters deleted. All data in subsequent columns are summaries of phenotypic information for mice homozygous for the respective TAD boundary deletions. These include effects on viability (fully lethal in red cross mark, subviable shown in red or yellow arrow to indicate magnitude of effect, no deviation from expected Mendelian ratios shown in green check marks), fertility of both male and female homozygous animals (n=3 each, green check marks indicate fertile animals), and overt physiological phenotypes observed. Column for chromatin change indicates presence of significant DI changes and/or significant alterations to intra-domain chromatin contact frequencies in the flanking TADs, assessed by Hi-C. The last column indicates whether significant expression changes were observed for genes in the vicinity of the deleted TAD boundary. n.a.=not applicable, n.d.=not determined, n.s.= not significant.

Our findings also have important implications for interpreting human whole genome sequencing data in clinical genetic settings. Position effects ensuing from large structural variations are well known in human genetics^41^ but it is often difficult to disconnect such deletions and/or rearrangements of functional sequences, such as protein-coding genes and enhancers, from effects that are due to removal of boundary sequences. It is worth noting that of the eight regions deleted from the mouse genes, none are completely deleted in ~760,000 available human genomes, consistent with their functional importance (gnomAD-SV v2.1^42^ and rCNV2 (https://doi.org/10.1101/2021.01.26.21250098). A critical future goal will be to determine the relationship between human mutations within the thousands of genome-wide TAD boundaries and their impact on human phenotypic variation and disease. The present study indicates that removal of the relatively small TAD boundary sequences themselves commonly causes molecular or organismal phenotypes and, therefore, structural variants that include TAD boundary deletions in human patients should be considered as potential causes of pathogenicity.

## Acknowledgements

This work was supported by U.S. National Institutes of Health (NIH) grants to L.A.P. and A.V. (UM1HG009421). Research was conducted at the E.O. Lawrence Berkeley National Laboratory and performed under U.S. Department of Energy Contract DE-AC02-05CH11231, University of California. Phenotyping performed by the UC Davis Mouse Biology Program (MBP) (www.mousebiology.org) was funded by an NIH administrative supplement to the KOMP2 grant, 3UM1OD023221-07S1, for phenotyping non-coding elements. We thank Renee Araiza and Louise Lanoue for assistance with coordinating work and analyzing phenotypes at the UC Davis MBP Program, and Michal Kosicki at LBL for help with bioinformatics.

## Author Contributions

S.R., I.B., D.E.D., A.V., and L.A.P. designed the study. S.R. and I.B. devised the prioritization strategy for identifying TAD boundaries for experimental deletion, with assistance from Y.F.Y. S.R., Y.Zhu, R.H., A.A., Q.P., A.N.H., J.G., E.M.M., K.V.M., I.P.-F., C.S.N., V.A., and S.T. performed experiments, including generating and characterizing knock-out mice. K.C.K.L. oversaw the KOMP2 phenotyping effort at the UC Davis Mouse Biology Program. S.R., Y.Zhang, B.L., G.K., M.B., M.T. analyzed data. S.R., D.E.D., A.V., and L.A.P. wrote the manuscript with input from the remaining authors.

## Methods

### ENCODE ChIP Seq data analysis and prioritization of TAD boundary deletion loci

Over 3300 genome-wide annotated TAD boundaries^5,10^ were ranked on a weighted score (see **Extended Data Figure 1**), encompassing strength of insulation based on CCCTC-binding factor (CTCF) occupancy, co-occupancy of subunits of the cohesin complex and the transcription factor Znf143^31^ and CTCF-binding conservation at orthologous regions in four different mammalian species^32^ (**Figure 1, Extended Data Figure 1**). We analyzed approximately 100 individual ChIP-Seq datasets to this effect (listed in **Supplementary Table 1**). CTCF-bound sites genome-wide were spatially clustered and individually scored based on the criteria highlighted above within regions 40 kb upstream and downstream of each TAD boundary. An overall score for each boundary was devised based on the intervening bound CTCFs, excluding boundaries overlapping protein-coding genes, thus enabling unambiguous interpretation of the functional necessity of TAD boundaries in mammalian development. Furthermore, boundaries where flanking TADs harbored genes encoding transcription factors important for development and preferentially (to the extent possible) showing a divergent pattern of tissue-specific expression were prioritized for *in vivo* deletion. Our selection criteria did not factor in the directionality of CTCF motifs when selecting TAD boundary loci for deletion (**Extended Data Figure 1)**.

### Experimental design *in vivo*

All animal work was reviewed and approved by the Lawrence Berkeley National Laboratory Animal Welfare Committee. Mice were monitored daily for food and water intake, and animals were inspected weekly by the Chair of the Animal Welfare and Research Committee and the head of the animal facility in consultation with the veterinary staff. The LBNL ACF is accredited by the American Association for the Accreditation of Laboratory Animal Care (AAALAC). TAD boundary knockouts were performed in *Mus musculus* FVB strain mice. Mice across developmental stages from embryonic day 10.5 through P0, as well as mice between weeks 4-16 were used in this study. Animals of both sexes were used in the analysis. Sample size selection and randomization strategies are included in individual method sections. Unless otherwise stated, all phenotyped mice described in the paper resulted from crossing heterozygous TAD boundary deletion mice together to allow for the comparison of matched littermates of different genotypes. Samples were dissected and processed blind to genotype where applicable.

### Generation of TAD boundary deletion mice

Transgenic mice were generated using the *Mus musculus* FVB strain and a CRISPR/Cas9 microinjection protocol as described before^43^. Briefly, Cas9 protein (Integrated DNA Technologies Cat. No. 1081058) at a final concentration of 20 ng/ul was mixed with sgRNA targeting the intended locus (50 ng/ul, for all sgRNAs combined), in microinjection buffer (10mM Tris, pH 7.5; 0.1 mM EDTA). The mix was injected into the pronuclei of single cell stage fertilized FVB embryos obtained from the oviducts of super-ovulated 7-8 weeks old FVB females mated to FVB males (See **Supplementary Table 6** for sgRNA sequences). The injected embryos were then cultured in M16 medium supplemented with amino acids at 37°C under 5% CO_2_ for ~2 hours. The embryos were subsequently transferred into the uteri of pseudo-pregnant CD-1 surrogate mothers. Founder (F0) mice were genotyped using PCR with High Fidelity Platinum Taq Polymerase (Thermo Fisher) to identify those with the desired NHEJ-generated deletion breakpoints. Sanger sequencing was used to identify and confirm deletion breakpoints in F0 and F1 mice **(Extended Data Figure 10, Supplementary Table 6** for CRISPR sgRNA templates and **Supplementary Table 7** for primer sequences and PCR amplicons). Between one and four F0 founders were obtained for each of the TAD boundary deletion loci, each of which were simultaneously assayed for possible inversions by PCR. Only those F0 founders that harbored clean deletion alleles were backcrossed to wild-type mice and bred to procure F1 heterozygous mice. Given that each of the deletions across founders were consistent in the NHEJ-mediated deletion span, only one founder line for each locus was eventually selected to expand breeding for experiments in this paper. Additional confirmation and visualization of the deleted TAD boundaries is evident in Hi-C contact matrices resulting from Hi-C experiments on tissue from homozygous TAD boundary mutants compared to wild-type mice (**Extended Data Figure 10**). The procedures for generating transgenic and engineered mice were reviewed and approved by the Lawrence Berkeley National Laboratory (LBNL) Animal Welfare and Research Committee. The described mouse lines are made available through the Mutant Mouse Resource and Research Center, www.mmrrc.org, and can be found in the MMRRC catalog using the regulatory region symbol, or the Research Resource Identifiers (RRID) (**Supplementary Table 8**).

### Assessment of Mendelian segregation and viability

Sample sizes were selected empirically based on our previous studies^44,45^. Mendelian segregation was initially assessed postnatally on animals resulting from heterozygous crosses, thus allowing for comparison of matched littermates of different genotypes. Where applicable, Mendelian ratios were assessed in embryological time points as necessitated by the phenotype on a case-by-case basis.

### *In situ* Hi-C library generation

Hi-C experiments were performed on *ex vivo* liver tissue from male mice at post-natal day 56. Upon euthanasia, liver samples were harvested, flash frozen in liquid nitrogen and pulverized before 1% formaldehyde cross-linking for 15 mins. Thawed crosslinked tissue was dissociated by a gentleMACS Tissue Dissociator using the factory-set program and filtered through a 40 μm BD-cellstrainer. Cell pellets were centrifuged at 1000g for 6 mins at 4 °C and overlaid with 3 ml 1M sucrose. The suspension was centrifuged at 2500g for 6 min at 4 °C. Pelleted nuclei were resuspended in 50 μL 0.5% SDS and incubated for 10 min at 62°C. SDS was quenched by adding Triton X-100 and incubation for 15 mins at 37 °C. Chromatin was digested using MboI (100U; NEB) at 37°C overnight with shaking (1000rpm). The enzyme was inactivated by heating 20 min at 62 °C. Fragmented ends were labeled with biotin-14-dATP (Life Technologies) using Klenow DNA polymerase (0.8 U μl^-1^; NEB) for 60min at 37 °C with rotation (900rpm). Ends were subsequently ligated for 4h at room temperature using T4 DNA Ligase (4,000 units; NEB). Reverse crosslinking was performed using Proteinase K (1mg, NEB) and incubation at 55 °C overnight. The digestion efficiency and ligation efficiency were checked by gel electrophoresis. Next, DNA was purified by using ethanol precipitation and sheared using a Covaris focused-ultrasonicator (M220; duty cycle: 10%; Power: 50, Cycles/burst: 200, Time: 70 seconds). After size selection and purification using SPRI beads (Beckman Coulter), DNA was biotin pulled-down using Dynabeads MyOne Streptavidin T1 beads (Life Technologies). Finally, sequencing libraries were prepared on T1 magnetic beads, and final PCR amplification was performed for seven cycles based on qPCR analysis. Bead-purified libraries were quantified with a Qubit and then diluted for size distribution assessment using High Sensitivity D1000 ScreenTape on a TapeStation (Agilent).

Hi-C was performed on two biological replicates each for both mutant and control samples for each of the TAD boundary deletion loci. We were unable to generate Hi-C data from tissue samples for TAD boundary deletion locus B1 given (i) the early embryonic lethality of homozygous mutants for this mouse line, (ii) the requirement of large amount of tissue for performing Hi-C experiments, (iii) limitations of using heterozygous mutants for Hi-C experiments, as these would make allele-specific downstream analyses problematic given the isogenic nature of our experimental mice.

### Hi-C data analysis

The Hi-C data processing pipeline is available at https://github.com/ren-lab/hic-pipeline. Briefly, Hi-C reads were aligned to the mouse mm10 reference genome using BWA-MEM ^46^ for each read separately, and then paired. For chimeric reads, only 5’ end-mapped locations were kept. Duplicated read pairs mapped to the same location were removed to leave only one unique read pair. The output bam files were transformed into juicer file format for visualization in Juicebox^47,48^. Contact matrices were presented as observed/expected contacts at 25-kb resolution and normalized using the Knight–Ruiz matrix balancing method^49^. Directionality index for each sample was also generated at 25-kb resolution as described previously^5^. Insulation score for each sample was generated at 25-kb resolution with 500-kb square as described previously^50^. For data in **Fig. 3**, Directionality Index (DI) scores of five bins on the right and five bins on the left were averaged, prior to calculating the difference. A higher DI delta score indicates a stronger boundary. A Wilcoxon rank-sum test was performed between KO and WT samples using DI delta scores, and a p-value ≤ 0.05 considered significant. As a negative control, the same statistical test was performed on ~2,900 TAD boundaries that do not overlap with deletions, and differences between WT and KO samples were assessed by the same statistical test, using a significance threshold of p ≤ 0.05.

We did not observe any other boundaries genome-wide that showed a significant difference in DI delta score that exceeded that of deleted boundaries B2, B3, B4, B6, and B8, for which we had observed significant changes in DI delta score. Twelve boundaries (0.4%) genome-wide showed changes that were significant (p≤0.05) and were quantitatively the same or exceeded those observed at deleted TAD boundaries B5 and B7, for which we did not observe significant changes in DI delta score (p>0.05; **Figure 3** and **Extended Data Figure 5**. Interaction frequencies between genomic loci were additionally visualized on Juicebox (**Extended Data Figure 5**) ^47,48^.

### RNA-seq and quantitative real time PCR

A panel of tissues including forebrain, midbrain, hindbrain, face, heart, upper and lower limbs and neural tube was collected in a standardized manner at E11.5 from homozygous mutants as well as littermate wild-type embryos for each of the TAD boundary deletion loci^34^. Samples were suspended in 100 μl of commercially available (Qiagen) RLT buffer. Total RNAs were isolated by using the Qiagen RNeasy Mini Kit (Cat. No. 74104). A set of two relevant tissue types was further selected for each of the TAD Boundary deletion loci B2-B8 and processed for RNA-seq libraries in a standardized manner. Sequencing libraries were prepared by Novogene, and sequenced on an Illumina NovaSeq6000 (150bp, paired-end). RNA-seq data was analyzed using the ENCODE Uniform Processing Pipelines (https://www.encodeproject.org/pipelines/) implemented at DNAnexus (https://www.dnanexus.com). Using the ENCODE RNA-seq (Long) Pipeline – 1 (single-end) replicate pipeline (code available from https://github.com/ENCODE-DCC/rna-seq-pipeline), reads were mapped to the mouse genome (mm10) using STAR align (V2.12). Genome wide coverage plots were generated using bam to signals (v2.2.1). Gene expression counts were generated for gencode M4 gene annotations using RSEM (v1.4.1). Differential expression analyses were performed by using the DESeq program in the R Statistical Package https://bioconductor.org/packages/3.3/bioc/vignettes/DESeq/inst/doc/DESeq.pdf ^51,52^. Statistically significant differentially expressed genes for relevant tissues for each TAD boundary deletion locus are listed in **Supplementary Table 4**. RNA-seq experiments were performed on two biological replicates each for homozygous mutants, as well as wild-type controls.

Quantitative PCR analysis of key developmental genes in the vicinity of each TAD boundary in a larger panel of E11.5 tissues did not identify any additional significant changes in gene expression (**Extended Data Figure 6).** For the comprehensive panel of tissues collected, RNA was isolated as described above and cDNA was synthesized using Omniscript RT (Qiagen Cat. No.205111) per standard methods. qPCR assays were performed for at least two genes, each in TADs immediately flanking the deleted boundary. Taqman Assay reagents (Life Technologies) were used for all targets including genes that were used to normalize expression levels. Taqman assays (Roche Applied Science) with gene-specific primer sequences were generated using the manufacturer’s online algorithm and are listed in (**Supplementary Table 9**). All amplicons span exon-exon junctions in order to prevent amplification of genomic DNA. 30μl assays dispensed in TaqMan Universal PCR 2X master mix (Applied Biosystems) were performed on LightCycler 480 (Roche) according to manufacturer’s instructions. All Ct values were manually checked. Relative gene expression levels were calculated using the 2^-ΔΔCT^ method^53^ normalized to the *Actb* housekeeping gene, and the mean of wild-type control samples was set to 1. At least three independent mutant samples and littermate and stage matched controls were assessed for each genotype/condition. We did not observe significant expression changes near deleted boundaries B3, B5 and B7. Considering RNA-seq analysis was performed on bulk tissue from a single developmental timepoint, we cannot exclude that expression changes are restricted to subsets of cells present in these tissues or may be more pronounced at other developmental timepoints.

### Comprehensive mouse phenotyping

Directly relevant results are summarized in **Figure 4**. Mutants and wild-type controls for four out of eight TAD boundary deletion loci (B3, B5, B6 and B7) underwent comprehensive phenotyping using a standardized pipeline at the Mouse Biology Program (MBP), University of California, Davis. The pipeline is part of the NIH-funded Knockout Mouse Phenotyping Project (KOMP2), a participant in the International Mouse Phenotyping Consortium (IMPC). Phenotyping tests are derived from the International Mouse Phenotyping Resource of Standardized Screens (IMPReSS), https://www.mousephenotype.org/impress ^54^, all protocols and metadata are accessible at https://www.mousephenotype.org/impress/PipelineInfo. The KOMP2 phenotyping (**Extended Data Figure 7**) and statistical analysis methods are previously described^55,56^. **Supplementary Table 5** summarizes other statistically significant (p<0.005) results reported. Mutants and wild-type controls for the other three TAD boundary deletion loci (B2, B4 and B8) were phenotyped using standard methods for gross necropsy, organ weights and histopathology for all major organ systems at the Comparative Pathology Laboratory, University of California, Davis.

### *In vivo* transgenic enhancer-reporter assay

Transgenic enhancer-reporter assays for the predicted lung enhancer (852bp) were performed in a site-directed transgenic mouse assay using a minimal *Shh* promoter and *lacZ* reporter gene (**Extended Data Figure 10**) at a non-endogenous, safe harbor locus ^43^. The predicted enhancer region was PCR amplified from mouse genomic DNA; chr5:120101603-120102454 (mm10), CTGGGCTACAGGAAGTTGGA (forward primer), CAGAGGGCATGAGAGAGACC (reverse primer), 852 bp PCR amplicon. The PCR amplicon was cloned into a *lacZ* reporter vector (Addgene #139098) using Gibson assembly (New England Biolabs)^57^. The final transgenic vector consists of the predicted enhancer–promoter–reporter sequence flanked by homology arms intended for site-specific integration into the *H11* locus in the mouse genome^43^. Sequence of the cloned constructs was confirmed with Sanger sequencing as well as MiSeq. Transgenic mice were generated using pronuclear injection, as described above for generating the TAD boundary deletion mice. F0 embryos were collected for staining at E11.5, E14.5 and E16.5.

b-galactosidase staining was performed as previously described^43^. Briefly, embryos were washed in cold 1× phosphate-buffered saline (PBS) and fixed with 4% paraformaldehyde (PFA) for 30 minutes for E11.5 and 60 minutes for E14.5 and E16.5 embryos, respectively, while rolling at room temperature. The embryos were washed in embryo wash buffer (2 mM magnesium chloride [Ambion, catalog no. AM9530], 0.02% NP-40 substitute [Fluka, catalog no. 74385], 0.01% deoxycholate [Sigma-Aldrich, catalog no. D6750] diluted in 0.1 M phosphate buffer, pH 7.3) three times for 30 minutes each at room temperature and transferred into freshly made X-gal staining solution (4 mM potassium ferricyanide [Sigma-Aldrich, catalog no. P3667], 4 mM potassium ferrocyanide [Sigma-Aldrich, catalog no. P9387], 20 mM Tris, pH 7.5 [Invitrogen, catalog no. 15567027], 1 mg ml^-1^ of X-gal [Sigma-Aldrich, catalog no. B4252]). Embryos were incubated in the staining solution overnight while rolling at room temperature and protected from light. Embryos were then washed with 1× PBS three times for 30-60 minutes per wash and subsequently stored in 4% PFA at 4 °C. The embryos were genotyped for presence of the transgenic construct as previously described^43^. Only those embryos positive for transgene integration into the *H11* locus and at the correct developmental stage were considered for comparative reporter gene activity across the three constructs tested. The exact number of embryos are reported in **Extended Data Figure 8**.

### Statistical analysis

Statistical analyses are described in detail in the Methods sections above. Whenever a p-value is reported in the text, the statistical test is also indicated. All statistics were estimated, and plots were generated using the statistical computing environment R (www.r-project.org).

### Data availability

The Hi-C and RNA-seq data discussed in this publication have been deposited in NCBI’s Gene Expression Omnibus^58,59^ and are accessible through GEO Series accession number GSE172089 (Reviewer access token: yvutceuufhsprub). Additional data supporting the findings of this study are available from the corresponding authors upon reasonable request.

## Extended Data

**Extended Data Figure 1.**
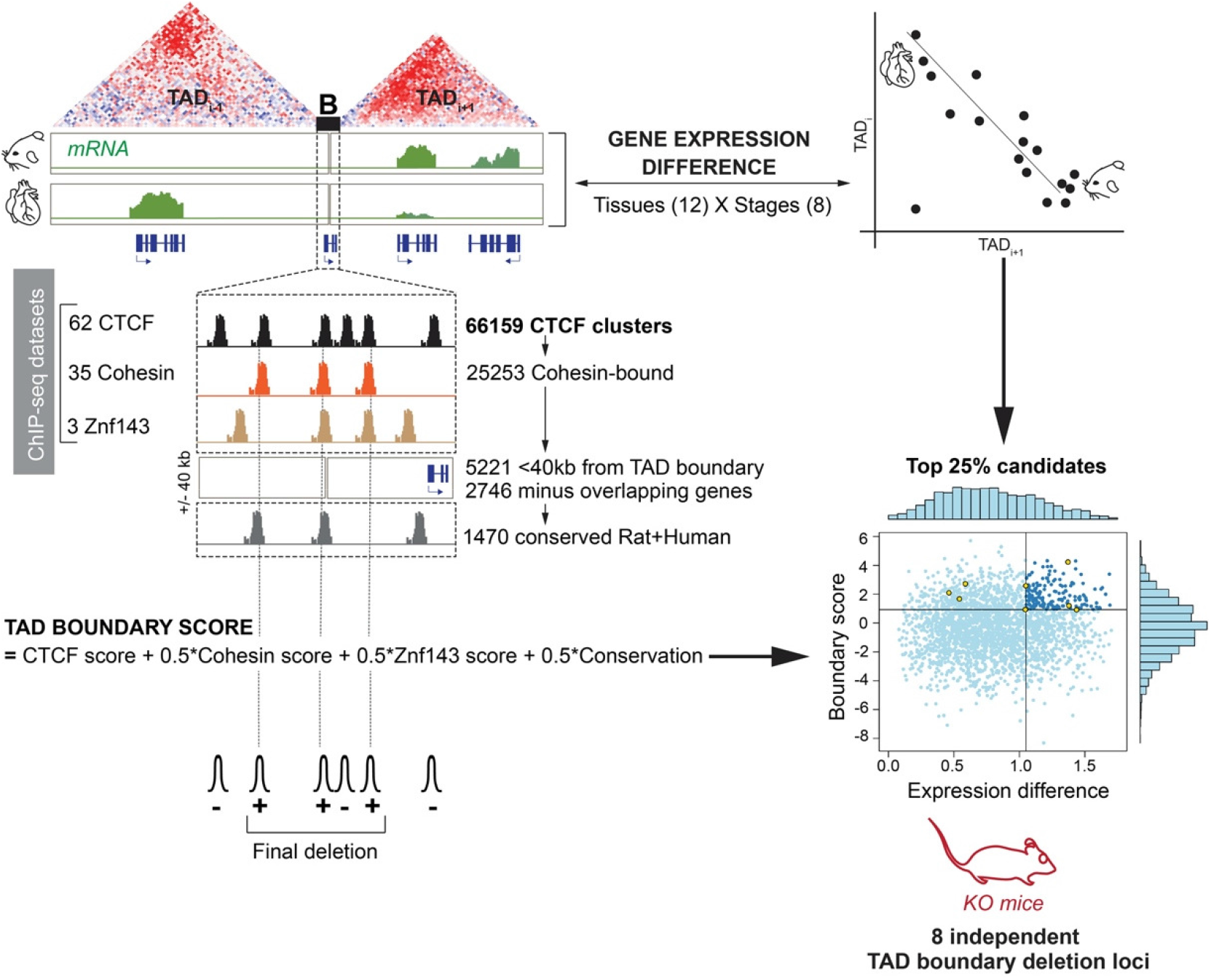
Schematic describing details of TAD boundary score and gene expression differences for selecting TAD boundary loci for deletion. **Left:** CTCF clusters at TAD boundaries were systematically scored for co-bound Cohesin and Znf14. Conservation of CTCF sites between rodent and human was also factored into the weighted score. **Top Right:** We used an extensive collection of gene expression data from embryonic mouse tissues to assess differences and similarities between developmental genes on opposite sides of each boundary (*He et al, Nature* 2020). The rationale for selecting TAD pairs harboring genes with divergent expression profiles was to enable straightforward scoring of molecular phenotypes that are expected to result from TAD boundary disruption and perturbation of typical contacts between *cis*-regulatory elements and their cognate genes. **Bottom right:** We selected 8 TAD boundaries for deletion (yellow points) from the top 25% of candidates based on high boundary score, as well as moderate to high gene expression differences for genes in neighboring TADs.

**Extended Data Figure 2.**
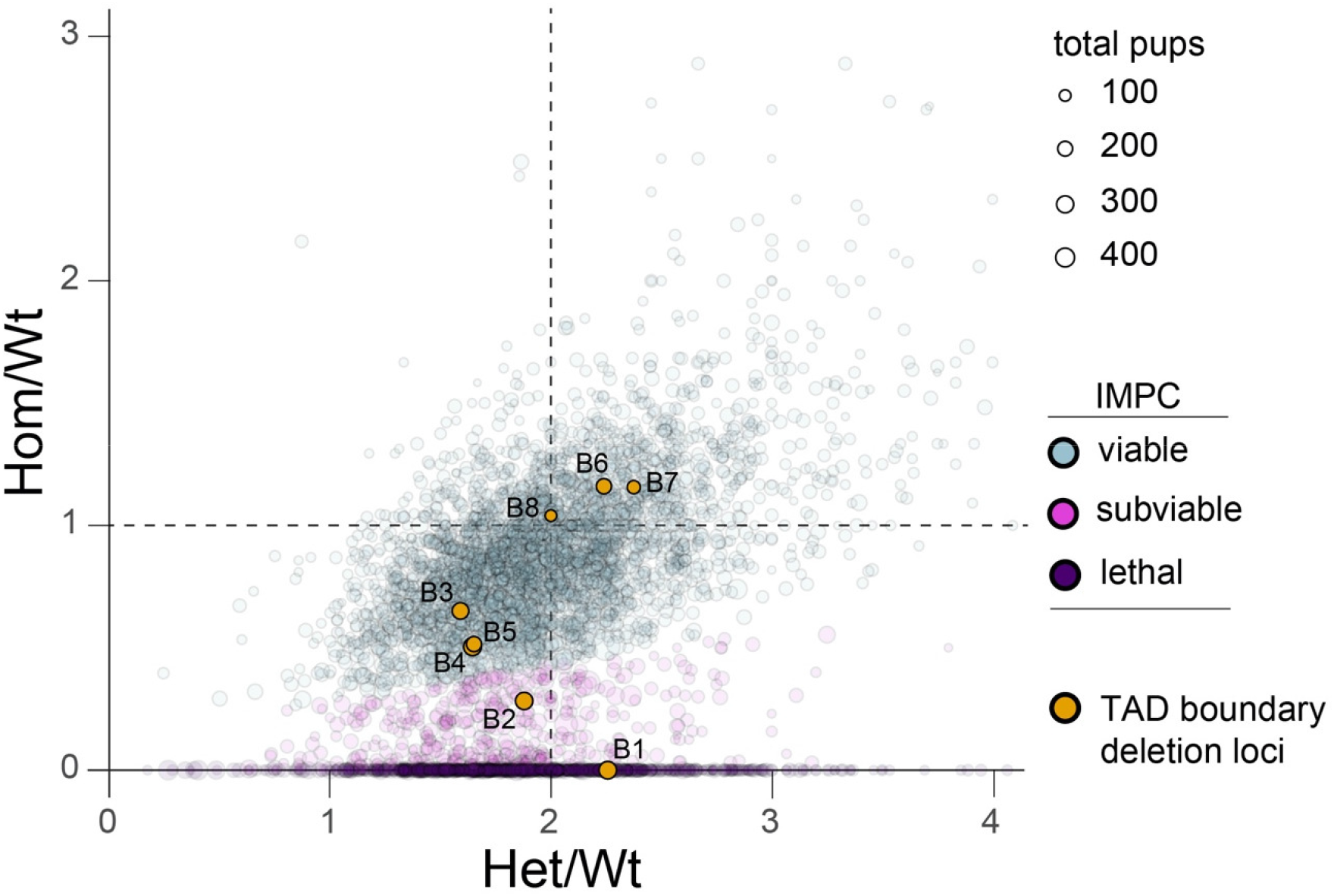
Allelic ratios of TAD boundary deletion KO lines compared with International Mouse Phenotyping Consortium (IMPC) mouse viability data. This comparison is based on data analyses from Cacheiro *et al*, Nat. Commun., 2020 and Mendelian segregation data publicly available for homozygous null mice generated by IMPC. This plot shows viability data by genotype ratios before weaning stages for independent KO lines corresponding to 4866 mouse genes from the IMPC curated data. Sizes of circles correspond to sample size of total pups genotyped for a given gene KO mouse line. IMPC defines categories as follows: KO lines are defined as “Viable” (gray) when homozygous, heterozygous and wild-type pups for a given gene KO are obtained at expected frequencies; KO lines where approximately half of the expected number of homozygotes (δ 12.5%) are obtained are defined as “subviable” (magenta); and KO lines where no homozygotes are obtained are defined as “lethal” (violet). The TAD boundary deletion lines in this study are shown in orange. No homozygotes were obtained at birth for locus B1, and boundary deletions for loci B2 and B3, respectively, resulted in approximately 65% and 20% fewer homozygous pups. B4 and B5 resulted in approximately 37% fewer homozygous pups at weaning. While this strongly suggests partially penetrant embryonic lethality in TAD boundary KO lines B3-5, deviations from the expected offspring ratios are within the range for lines commonly reported as viable in IMPC gene knockout analyses. See **Figure 3** and **Methods** for details.

**Extended Data Figure 3.**
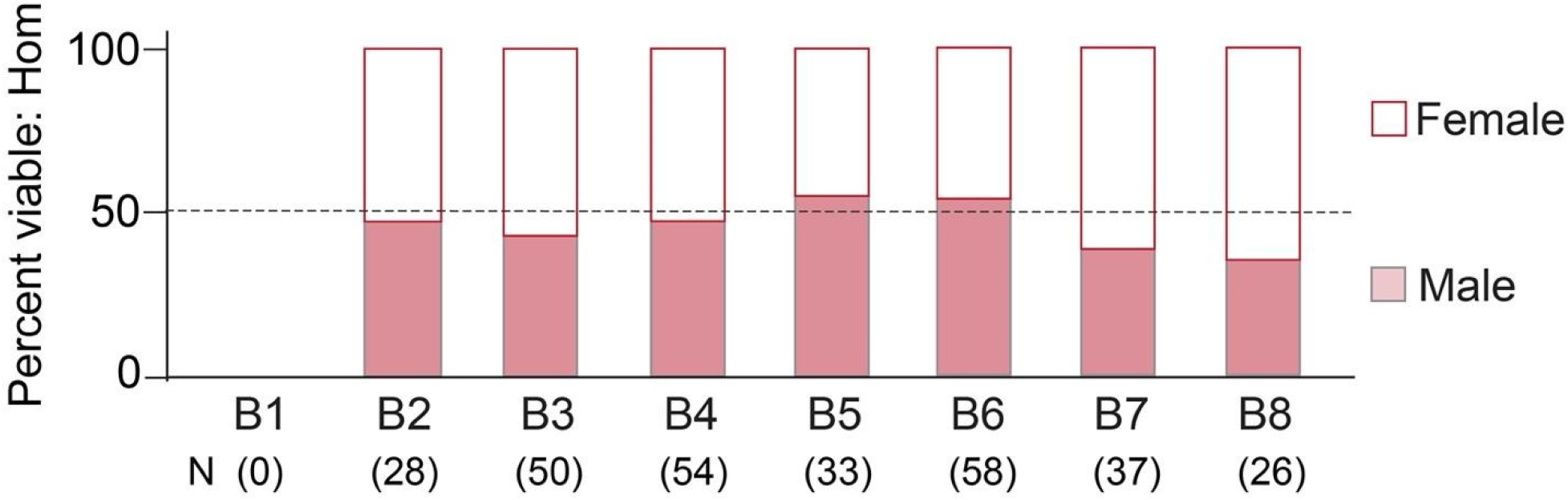
TAD boundary deletions show no sex bias within homozygous mutants at weaning. Mendelian segregation of male and female homozygous offspring from heterozygous crosses at weaning (approximately postnatal day 20, or P20) for all TAD boundary deletion lines.

**Extended Data Figure 4.**
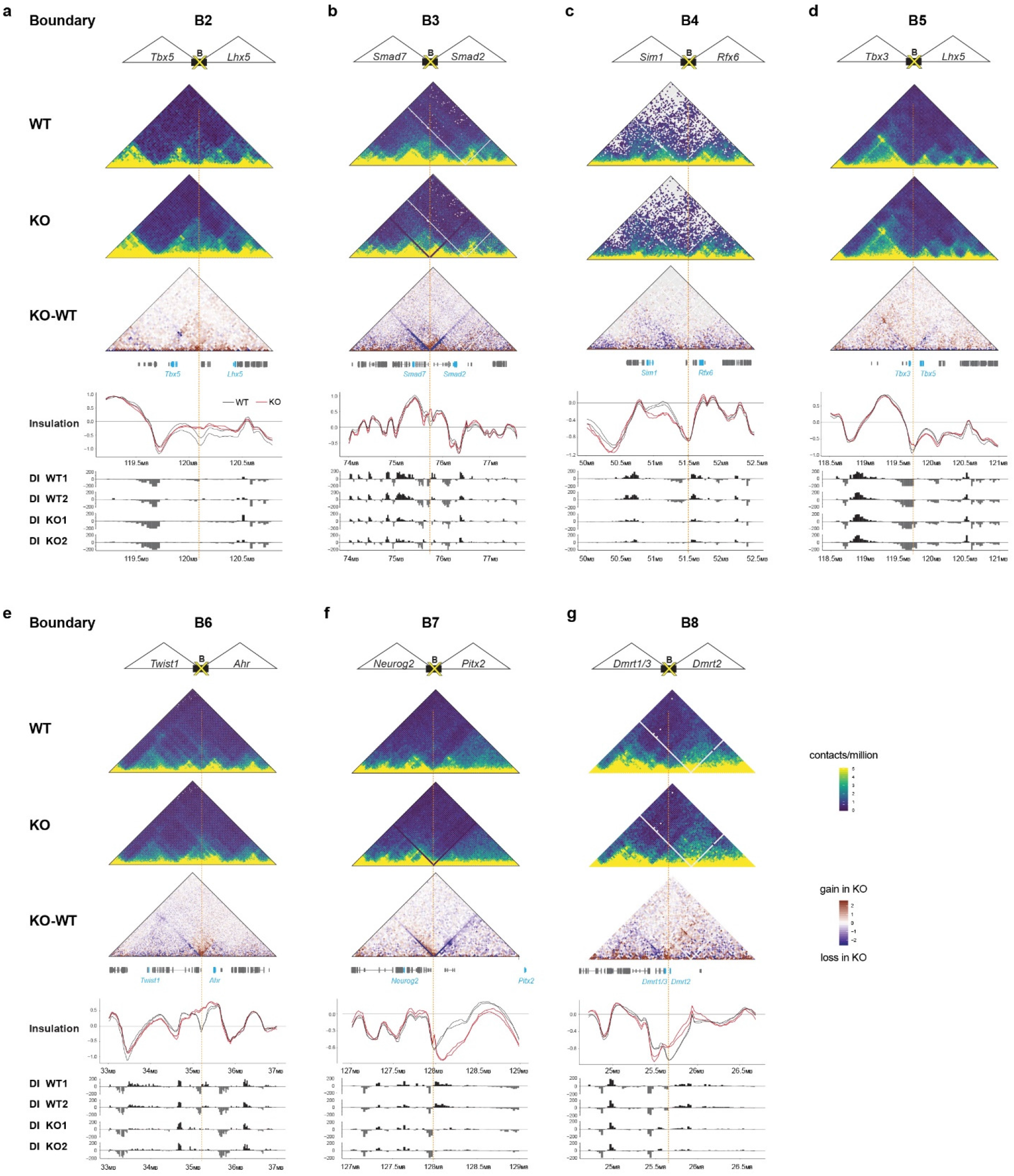
TAD boundary deletions result in abnormal TAD architecture. **a-g)** Top: Hi-C interaction maps of TAD boundary loci B2-8 showing interaction frequencies in representative wild-type (WT) and knockout (KO) mouse liver tissue. Bottom: Insulation profiles and Directionality Index (DI) for two wild-type and two homozygous mutants, i.e. two biological replicates for each of the respective TAD boundary KO lines is shown. See **Methods** for additional details.

**Extended Data Figure 5.**
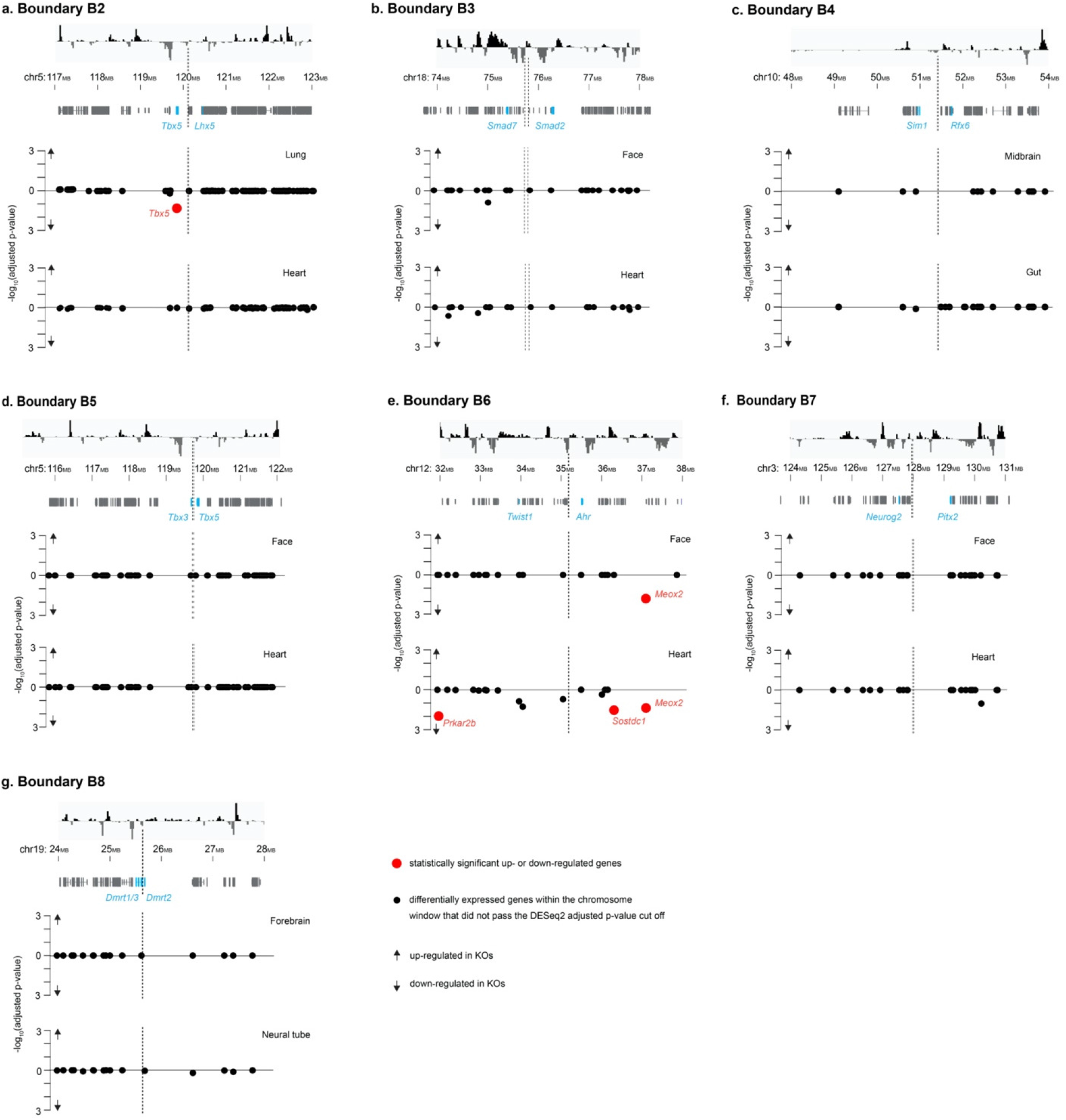
Gene expression changes in TAD boundary KOs. **a-g**) Local gene expression changes in two selected tissues at E11.5 for each of the TAD boundary deletion lines. Top panel shows Directionality Index track for reference, along with chromosomal coordinates below. Gray bars show RefSeq genes in the respective regions. Genes highlighted in blue are developmentally-expressed genes flanking the boundary. For gene expression plots in the lower panels, the x-axis denotes genomic location along the relevant chromosome, while the y-axis represents log10(adjusted p-value) and shows the specific tissues in which RNA-seq was performed. The vertical, dashed gray line indicates the position of the TAD boundary. Homozygous mutants (n=2) compared to wild-type (n=2) mice were used for each of the TAD boundary KO lines B3-8; an additional 2 samples (n=4) for each of the genotype groups were used for B2. Red points (**a,e**) indicate statistically significant up- or down-regulated genes (adjusted p-value < 0.05 using an FDR < 5%, see **Methods** for details).

**Extended Data Figure 6.**
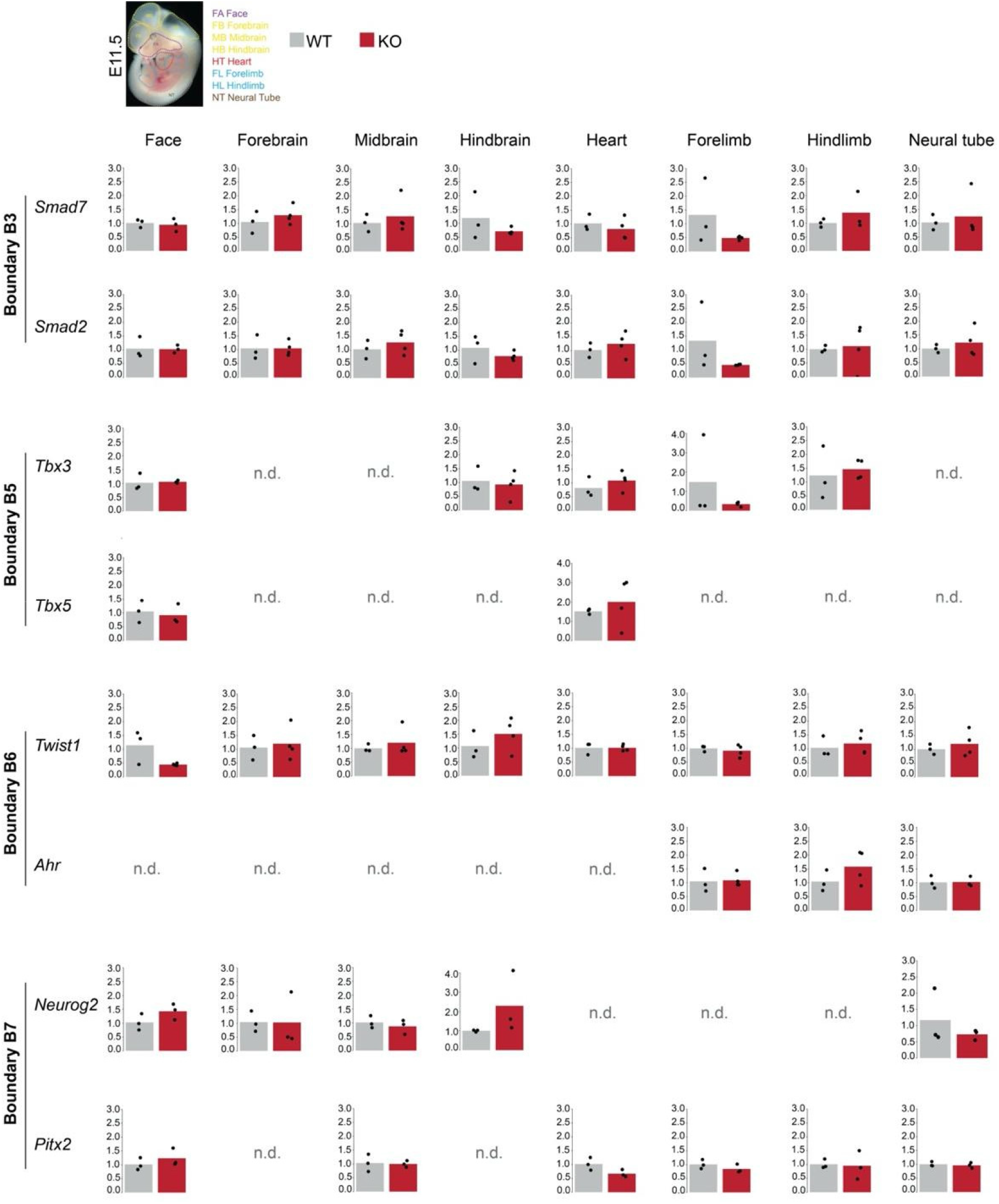
qPCR gene expression for select genes within flanking TADs. Quantitative real-time PCR results for TAD boundary KO lines B3 and B5-7 showing normalized gene expression (y-axis). qPCR results from a panel of tissues (face, forebrain, midbrain, hindbrain, heart, forelimb, hindlimb and neural tube) from homozygous mutants (KO, red bars) and wild-type (WT, black bars) embryos at E11.5 reveals lack of significantly up- or downregulated transcript levels in selected developmentally important genes, each present in a TAD adjacent to the deleted boundary. Bars denote average gene expression levels, while black dots show individual samples (n=3) for each of the KO and WT groups. p-value=0.05, from un unpaired, two-tailed *t*-test was considered as statistically significant. n.d.=not determined. Image on top shows a wild-type E11.5 mouse embryo for illustrative purpose outlining the tissue dissection intervals.

**Extended Data Figure 7.**
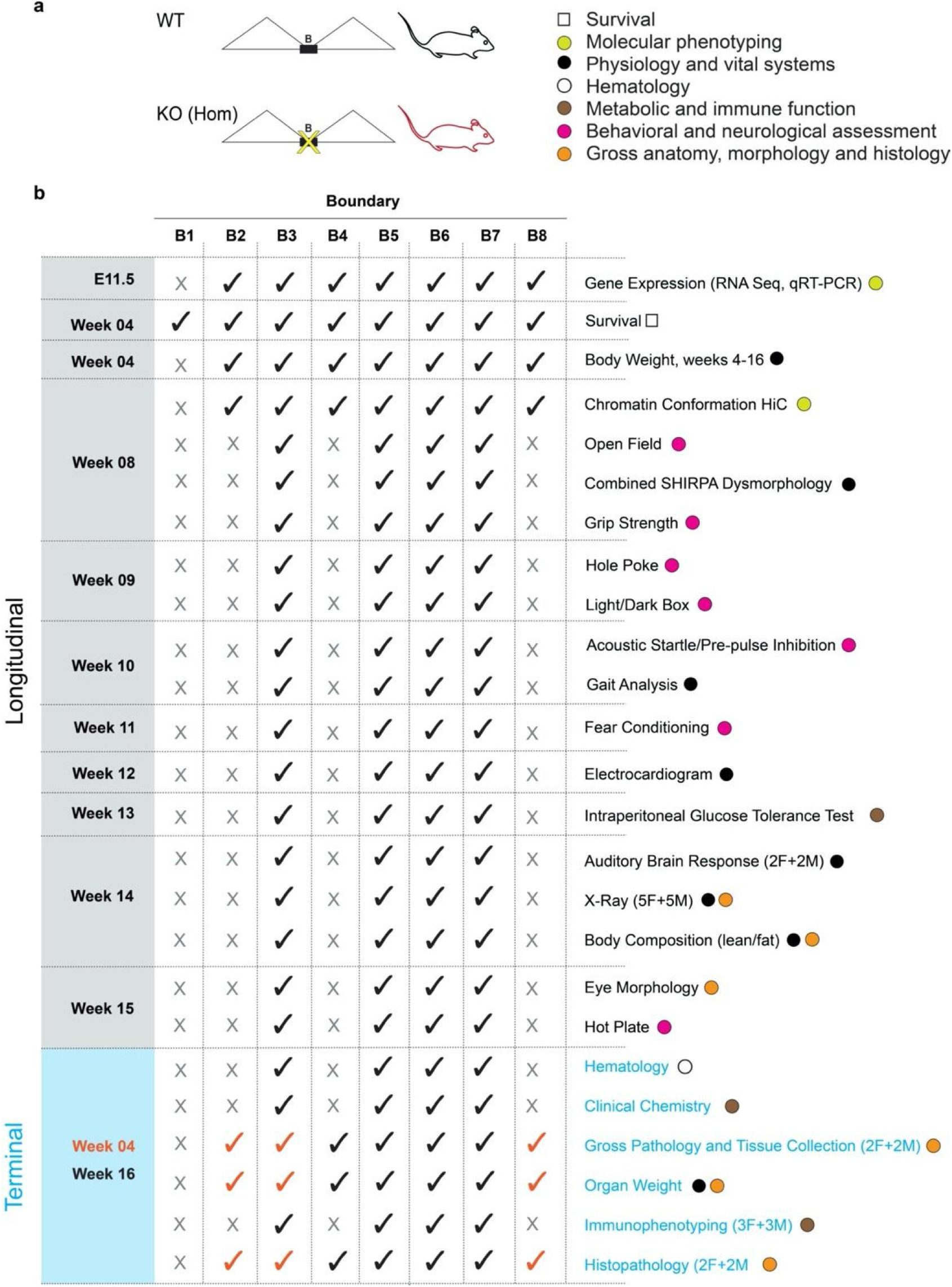
Comprehensive phenotyping strategy. **a)** An overview of the phenotyping strategy, including broader categories of phenotypic assessment and test types. **b)** Details of tests conducted. **Left**: shows age (embryonic or postnatal) at which the tests were performed in homozygous mutants and wild-type controls. **Middle:** Check marks indicate phenotyping test was performed on that line, while “X” indicates phenotyping test not performed. **Right**: The names of the tests listed using standard terminology. More details are described in the Methods section.

**Extended Data Figure 8.**
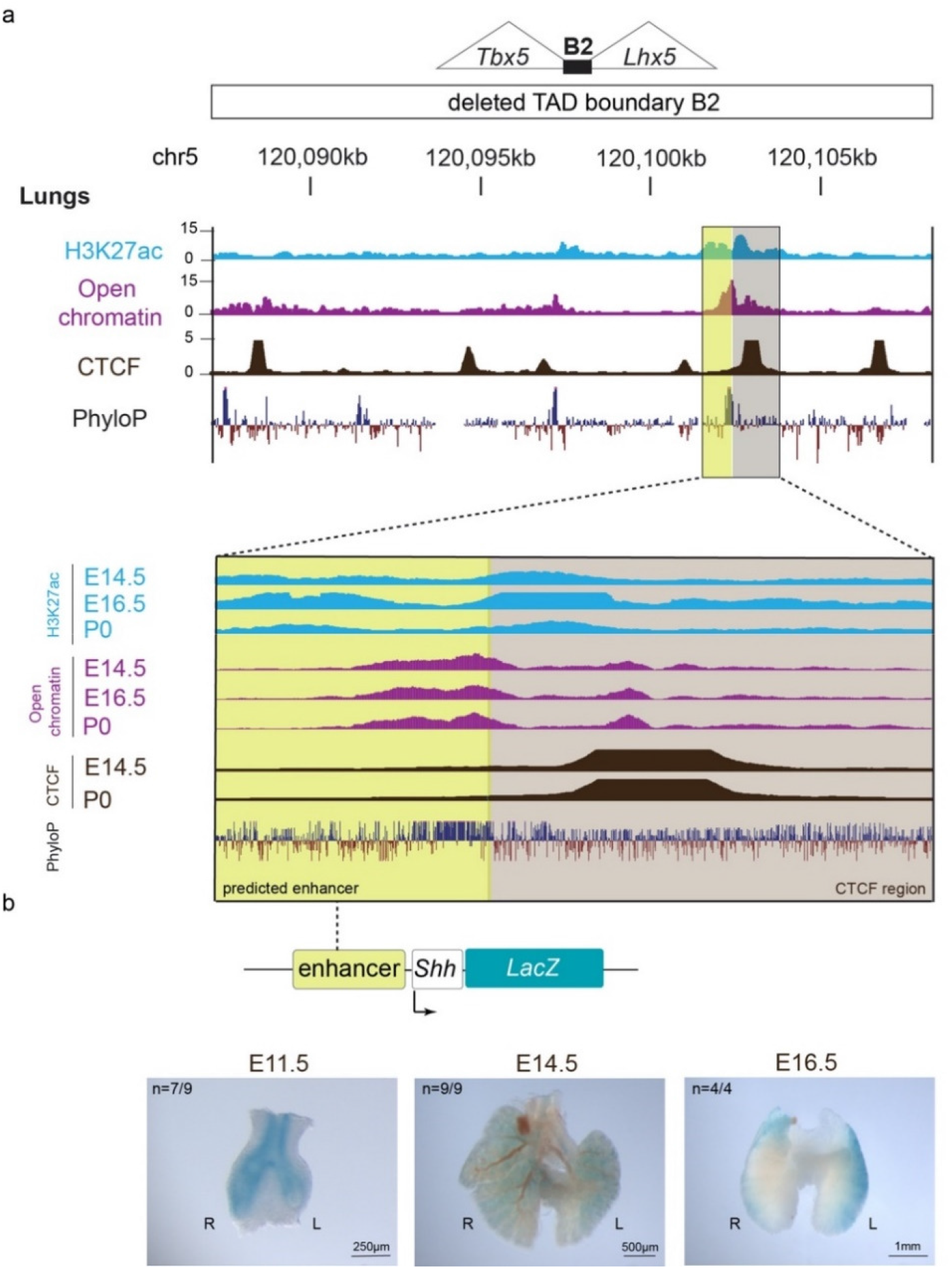
Regulatory element within TAD boundary B2 functions as lung-specific enhancer. **a)** Upper panel shows the 21 kb span of the TAD boundary B2, in the vicinity of *Tbx5*, with histone modification, open chromatin data from ATAC-seq and placental conservation (PhyloP scores) from publicly available ENCODE3 data and the UCSC Genome Browser. Tracks for CTCF-bound regions are from ChIP-seq data summarized in Table 1. All tracks shown are for mouse lung tissue. The lower panel shows a magnified view of the predicted lung enhancer (highlighted yellow, 852 bp), along with its adjacent insulating CTCF-bound region (highlighted beige, 1.3 kb). H3K27ac bound regions (blue), open chromatin regions from ATAC-seq assays (purple) and CTCF-bound regions (dark brown) are shown along with the respective mouse developmental stages indicated on the far left of the tracks. **b) Top:** Cartoon showing that the predicted enhancer was cloned upstream of the *Shh* promoter-*LacZ* for transgenic enhancer-reporter assay, described in detail in Methods. **Bottom:** Brightfield images of the lungs at E11.5, 14.5 and 16.5 respectively, for the predicted enhancer element driving positive β-galactosidase staining in the lungs at all stages examined. n, independent biological replicates with reproducible results out of the total transgenic embryos obtained. Scale bars, 250μm (E11.5), 500μm (E14.5), and 1mm (E16.5).

**Extended Data Figure 9.**
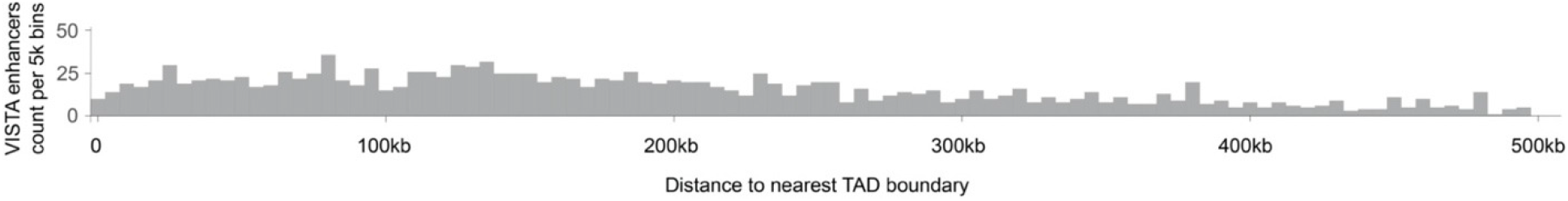
Enhancer-TAD boundary distances. Distances from nearest TAD boundary for over 1,800 unique enhancer elements from the VISTA Enhancer Browser that had validated enhancer activity *in vivo*. ~275 elements mapped ≥500 kb from the nearest TAD boundary. Bin size = 5 kb (*Zhan et al*, Genome Research 2017). The median enhancer-TAD boundary distance is 201 kb. An apparent mild depletion in enhancers in the immediate vicinity of boundaries may be due to biases in the selection of enhancers in the VISTA database. We observed multiple additional enhancers with validated *in vivo* activity during embryonic development that are located in the immediate (0-5kb) or extended vicinity of TAD boundaries.

**Extended Data Figure 10.**
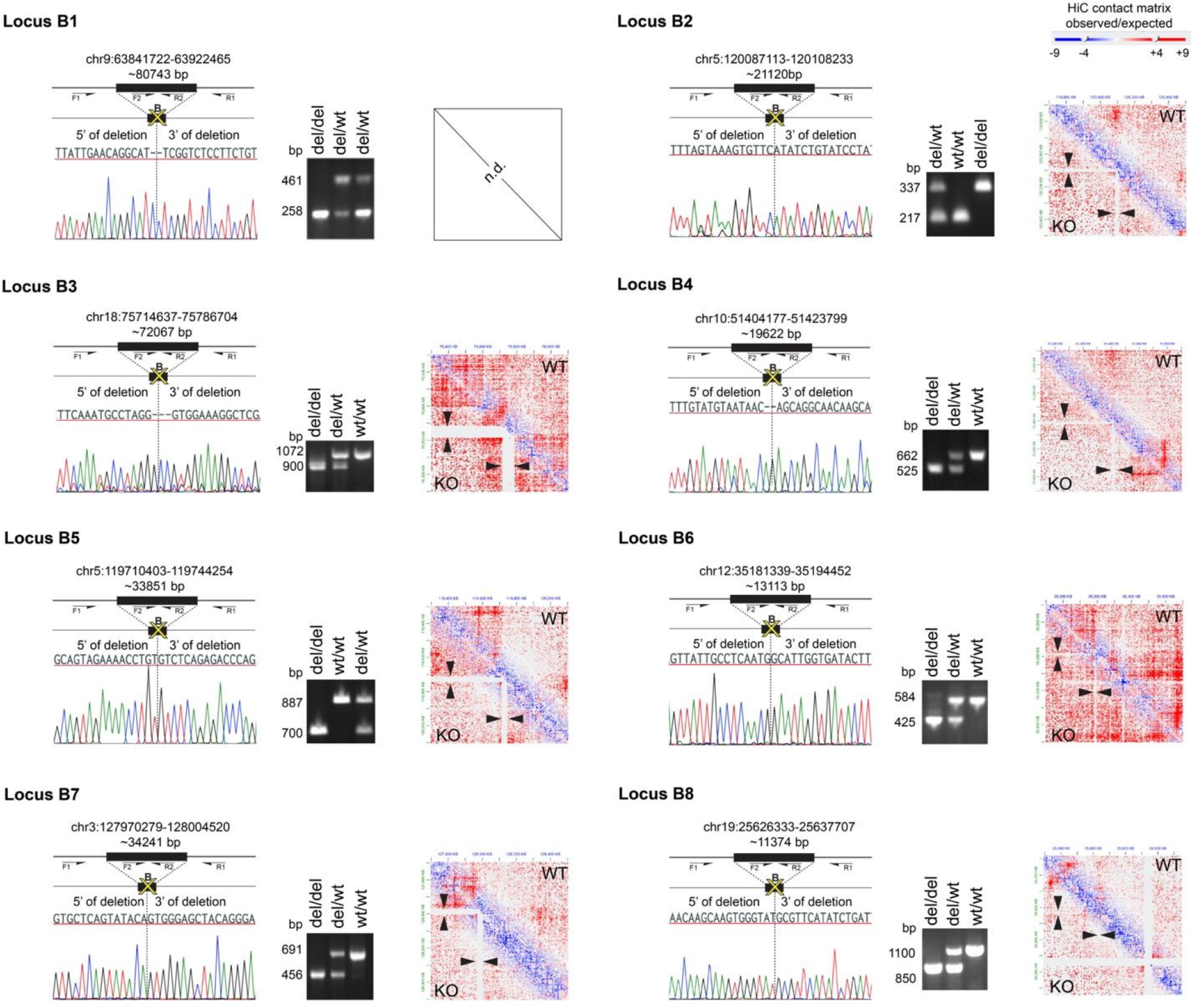
CRISPR-mediated deletion of eight TAD boundaries. For each of the TAD boundary deletion loci B1-8, left hand side panels show boundary regions (B) depicted as solid black bars, along with the chromosomal coordinates (mm10) and deletion span. Yellow crosses coincide with CRISPR-mediated deletion breakpoints. Below, Sanger sequencing traces show the deletion breakpoints (indicated by the vertical dashed line) for the TAD boundary deletion alleles. Black arrows indicate forward and reverse primers used for PCR validation. See Supplementary Tables 5 and 7 for sgRNA and PCR primer sequences. Middle panels show PCR genotyping and amplicon sizes for TAD boundary deletion (del/del), heterozygous (del/wt), and wild-type (wt/wt) animals. Right hand side panels show Hi-C interaction heatmaps (Juicebox output) of TADs from a wild-type and a representative TAD boundary knockout (KO) for each of the deletion lines, for the respective genomic regions. Black arrowheads indicate the region corresponding to the TAD boundary deletion in a homozygous mutant juxtaposed with the identical region from the wild-type control.

## Notes

### Competing Interest Statement

The authors have declared no competing interest.

